# Knockout of the LRRK2-counteracting RAB phosphatase PPM1H disrupts axonal autophagy and exacerbates alpha-synuclein aggregation

**DOI:** 10.1101/2024.10.14.618089

**Authors:** Michel Fricke, Anna Mechel, Björn Twellsieck, Jessica M. Grein, Maria-Sol Cima-Omori, Markus Zweckstetter, Erika L.F. Holzbaur, C. Alexander Boecker

## Abstract

Parkinson disease-causing mutations in the *LRRK2* gene hyperactivate LRRK2 kinase activity, leading to increased phosphorylation of a subset of RAB GTPases, which are master regulators of intracellular trafficking. In neurons, processive retrograde transport of autophagosomes is essential for autophagosome maturation and effective degradation of autophagosomal cargo in the axon. We found that knockout of the LRRK2-counteracting RAB phosphatase PPM1H resulted in a gene dose-dependent disruption of the axonal transport of autophagosomes, leading to impaired degradation of axonal alpha-synuclein (aSyn), a key protein in Parkinson disease pathophysiology. Defective autophagosome transport and impaired aSyn degradation also correlated with increased aSyn aggregation in primary *PPM1H* knockout neurons exposed to preformed fibrils of aSyn, an effect that was dependent on LRRK2 kinase activity. Thus, our results link LRRK2-mediated RAB hyperphosphorylation to aSyn pathology in Parkinson disease and further establish a role for impaired autophagy in Parkinson disease pathophysiology.

## INTRODUCTION

Pathogenic mutations in the leucine-rich repeat kinase 2 (*LRRK2*) gene cause Parkinson disease (PD). These mutations induce increased LRRK2 kinase activity, leading to hyperphosphorylation of a subset of RAB GTPases [1,2]. Increased LRRK2 kinase activity and RAB hyperphosphorylation are also induced indirectly by the PD mutations *VPS35*-p.D260N and *RAB32*-p.S71R [3–5] and have been observed upon exposure to PD-causing environmental toxins [6,7] as well as in idiopathic PD cases [8]. Furthermore, a recent report identified a PD case with a truncating mutation of protein phosphatase 1H (PPM1H), the main phosphatase counteracting LRRK2 kinase activity by specifically dephosphorylating RAB proteins targeted by LRRK2 [9,10]. Taken together, these observations suggest that different causes of PD may converge on a similar core mechanism of elevated LRRK2-mediated RAB phosphorylation. However, it is still unclear how RAB hyperphosphorylation, resulting from an imbalance between LRRK2 kinase and PPM1H phosphatase activity, mediates the neurodegeneration found in patients with PD.

In previous work, we found that LRRK2-mediated RAB hyperphosphorylation disrupts the axonal transport of autophagic vesicles (AVs) [11,12]. AVs are primarily formed in the distal axon and at presynaptic sites before undergoing processive dynein-driven retrograde transport toward the soma [13–16]. During retrograde transport, AVs fuse with lysosomes, lowering the intra-lumenal pH and activating degradative enzymes [17,18]. Importantly, processive AV transport is tightly linked to effective AV maturation and degradative function [19–21]. Hyperactive LRRK2 inappropriately activates competing kinesin motors on the AV, leading to an unregulated tug-of-war that disrupts transport and induces defective maturation of axonal AVs [11,12]. Together, these observations suggest that the hyperphosphorylation of RABs by mutated LRRK2 or resulting from loss of counteracting PPM1H phosphatase may impair efficient degradation of autophagosomal cargo [11,12].

Alpha-synuclein (aSyn) is an autophagosomal cargo of particular interest [22–24], as the aggregation of aSyn into Lewy bodies and Lewy neurites is the pathological hallmark of PD and a cardinal feature of PD disease progression. aSyn is predominantly found at presynaptic sites, the primary location of neuronal AV biogenesis, and aSyn aggregation to Lewy neurites has been shown to occur early during PD pathogenesis, preceding the formation of Lewy bodies in the neuronal soma [25]. Defective degradation of aSyn is predicted to contribute to the development of PD, as increased levels of endogenous aSyn correlate with increased aSyn aggregation upon treatment with preformed fibrils (PFFs), and *SNCA* gene duplications or triplications cause familial forms of PD [26–28]. Interestingly, previous work has shown increased formation of axonal aSyn aggregates following PFF treatment in different types of neurons expressing hyperactive LRRK2 [29–33]. However, it remains unclear how LRRK2 mutations augment aSyn pathology.

Here, we leveraged knockout (KO) models of the LRRK2 counteracting phosphatase PPM1H to investigate whether the effects of pathogenic LRRK2 on aSyn pathology are mediated by RAB hyperphosphorylation. We found that primary *PPM1H* KO neurons exhibited LRRK2 kinase activity-dependent defects in retrograde AV transport, which was accompanied by impaired autophagosomal degradation of axonal aSyn-EGFP. Furthermore, we found that *PPM1H* KO increased aSyn aggregation upon PFF treatment in primary neurons. This effect was rescued by pharmacological LRRK2 kinase inhibition, suggesting that increased aSyn aggregation is caused by LRRK2-mediated RAB hyperphosphorylation. Together, our data further link increased RAB phosphorylation to defects in axonal autophagy and demonstrate that LRRK2-mediated RAB hyperphosphorylation increases neuronal vulnerability to aSyn aggregation.

## RESULTS

### Knockout of the LRRK2-counteracting phosphatase PPM1H disrupts AV transport in mouse cortical neurons

KO of the LRRK2-counteracting RAB phosphatase PPM1H was previously determined to disrupt the axonal transport of autophagic vesicles (AVs) in human iPSC-derived *PPM1H^-/-^* neurons [12]. To assess whether this phenotype is consistent across different model systems and to test the effect of a heterozygous loss of PPM1H, we investigated the axonal transport of mScarlet-LC3B labeled AVs in *PPM1H*^+/+^, *PPM1H*^+/-^, and *PPM1H*^-/-^ primary neurons from *PPM1H* KO mice (Figure 1A) [34]. Western blots showed increased levels of LRRK2-phosphorylated pT73 Rab10 in PPM1H^-/-^ neurons and a trend toward increased pT73 Rab10 levels in *PPM1H^+/-^*neurons (Figure S1A-B). We found that decreased expression of PPM1H in primary neurons markedly affected the directionality of axonal AVs by increasing the stationary fraction of AVs at the expense of the retrograde fraction (Figure 1B). The increase in stationary AVs was significantly higher in *PPM1H^-/-^*than in *PPM1H^+/-^* neurons, suggesting a gene dose-dependent effect. To further characterize the effect of loss of PPM1H on AV transport, we quantified non-processive motility and pausing behavior of motile AVs. We found that both heterozygous and homozygous *PPM1H* KO resulted in reduced AV processivity compared to *PPM1H^+/+^*, with a higher number of directional reversals (Figure 1C) and increased Δ run length, a metric of non-processive motility calculated as the difference between total run length and net run length of each AV (Figure S1C). Furthermore, motile AVs in *PPM1H^+/-^*and *PPM1H^-/-^* neurons showed a significant increase in the number of pauses and in pause duration (Figure 1D and Figure S1D). This robust increase in non-processive motility and pausing is consistent with the model of an underlying “tug-of-war” between anterograde and retrograde motor proteins that was proposed in previous work [11,12]. Together, the increased pausing of motile AVs and the higher number of stationary AVs resulted in a marked increase in the overall fraction of time that *PPM1H* KO AVs spent paused (Figure 1E). Due to their even higher number of stationary AVs, the overall fraction of time paused was significantly longer in *PPM1H^-/-^* neurons than in *PPM1H^+/-^*neurons (Figure 1E).

**Figure 1.**
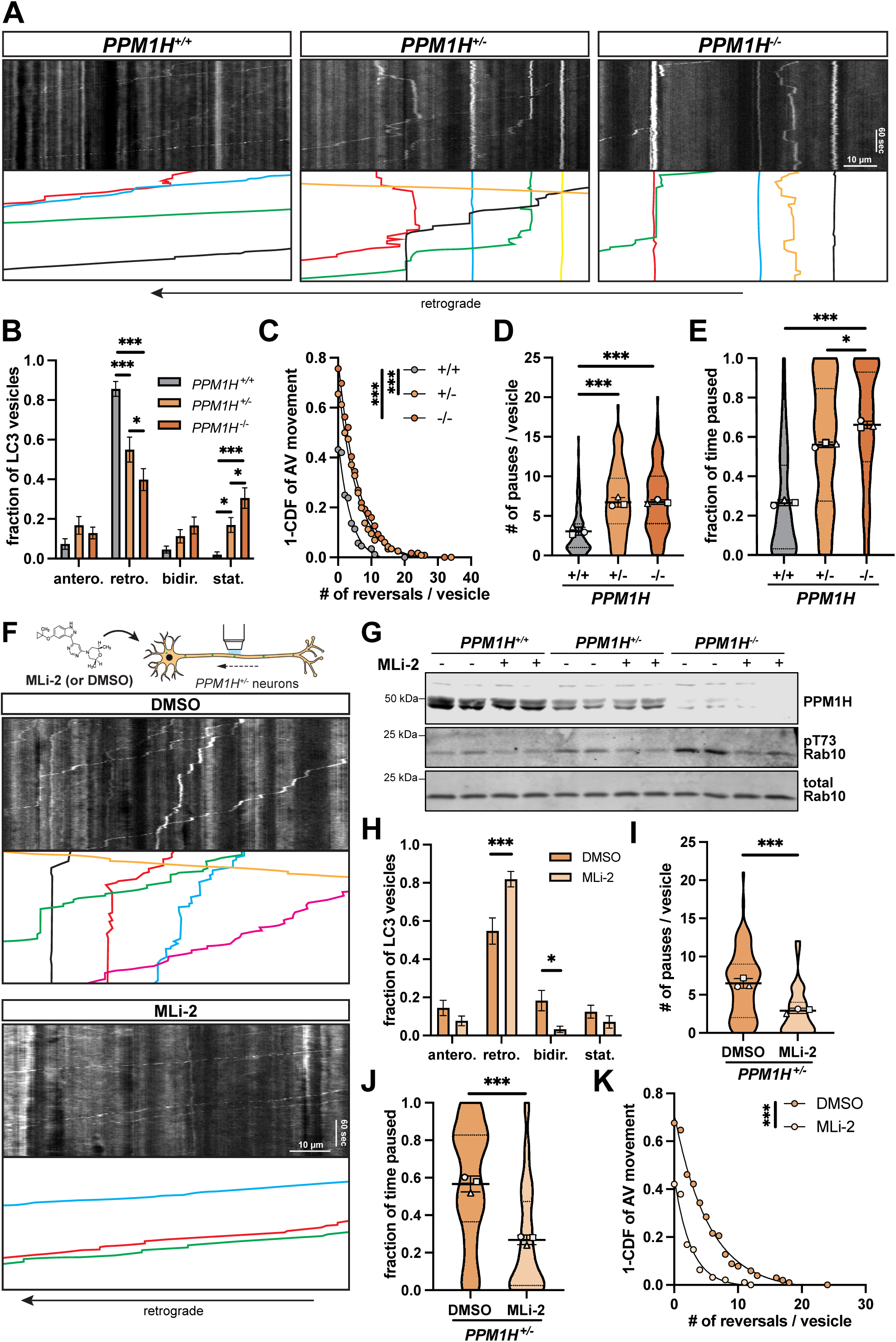
Knockout of PPM1H in primary neurons disrupts axonal AV transport in a LRRK2 kinase-dependent manner. (**A**) Kymographs of axonal mScarlet-LC3+ vesicles in *PPM1H^+/+^*, *PPM1H^+/-^*, and *PPM1H^-/-^* mouse cortical neurons. Example AV traces are highlighted. Scale bars, 10 µm (horizontal) and 60 sec (vertical). (**B**) Directionality of AVs in *PPM1H^+/+^*, *PPM1H^+/-^*, and *PPM1H^-/-^* neurons. Antero., anterograde; retro., retrograde; bidir., bi-directional; stat., stationary (mean ± SEM; n = 27-29 neurons from 3 independent experiments; *p<0.05; **p<0.01; ***p<0.001; two-way ANOVA with Sidak’s multiple comparisons test). (**C-D**) Directional reversals (C) and pause number (D) of motile AVs in *PPM1H^+/+^*, *PPM1H^+/-^*, and *PPM1H^-/-^* neurons (mean ± SD for panel D; n = 88-119 motile AVs from 27-29 neurons from 3 independent experiments; ***p<0.001; mixed effects model analysis, see Methods for specific models used). (**E**) Fraction of time paused of all AVs in *PPM1H^+/+^*, *PPM1H^+/-^*, and *PPM1H^-/-^* neurons (mean ± SD; n = 90-167 AVs from 27-29 neurons from 3 independent experiments; ***p<0.001; mixed effects model analysis). (**F**) Kymographs of axonal mScarlet-LC3+ vesicles in *PPM1H^+/-^* neurons treated overnight with DMSO or 100 nM MLi-2. (**G**) Western blot of PPM1H, pT73 Rab10, and total Rab10 in *PPM1H^+/+^, PPM1H^+/-^*, and *PPM1H^-/-^*mouse cortical neurons treated with DMSO or 100 nM MLi-2 overnight. (**H**) Directionality of AVs in *PPM1H^+/-^* neurons treated with DMSO or MLi-2 (mean ± SEM; n = 23-24 neurons from 3 independent experiments; *p<0.05 ; ***p<0.001; two-way ANOVA with Sidak’s multiple comparison test). (**I-K**) Pause number (I) of motile AVs, fraction of time paused (J) of all AVs, and directional reversals (K) of motile AVs in *PPM1H^+/-^* neurons treated with DMSO or MLi-2 (mean ± SD for panel I and J; n = 95-102 motile AVs (I, K) and 100-118 total AVs (J) from 23-24 neurons from 3 independent experiments; ***p<0.001; mixed effects model analysis). For panel D, E, I, and J, scatter plot points indicate the means of three independent experiments. For panels C and K, curve fits were generated using nonlinear regression (two phase decay).

To determine whether the observed transport defects are LRRK2-dependent, we treated primary *PPM1H* KO neurons with the selective LRRK2 kinase inhibitor MLi-2 (Figure 1F-G) [35]. We focused on heterozygous *PPM1H^+/-^* neurons, which showed significant impairment of AV transport and were more readily available than homozygous *PPM1H^-/-^* neurons from breeding heterozygous mice. Treatment with MLi-2 significantly increased the retrograde fraction of AVs in *PPM1H^+/-^* neurons (Figure 1H). MLi-2 furthermore decreased the number of pauses of motile AVs, restoring the overall fraction of time paused to levels similar to *PPM1H^+/+^* neurons (Figure 1I-J). Non-processive motility, as measured by number of reversals, was also rescued by LRRK2 inhibition (Figure 1K).

Together, these findings show that loss of PPM1H disrupts axonal AV transport in primary neurons in a LRRK2 kinase activity-dependent manner. Results from homozygous *PPM1H* KO neurons showed a more severe transport impairment than seen in heterozygous PPM1H KO neurons. This suggests that the magnitude of AV transport impairment scales with the level of RAB hyperphosphorylation by LRRK2.

### Impaired AV transport is associated with defective autophagosomal degradation of aSyn-EGFP

Aggregation of aSyn to Lewy bodies and Lewy neurites is the pathological hallmark of PD and a cardinal feature of disease progression. aSyn is an established substrate of autophagy and is predominantly found at presynaptic sites in the distal axon, the primary location of neuronal AV biogenesis [22–24]. Upon formation in the distal axon, AVs are transported retrogradely towards the cell soma, maturing *en route* by fusion with lysosomal vesicles (Figure 2A) [13–17]. Previous experiments have shown that defects in AV transport are associated with impaired maturation of axonal AVs during retrograde transport [11,19,21].

**Figure 2.**
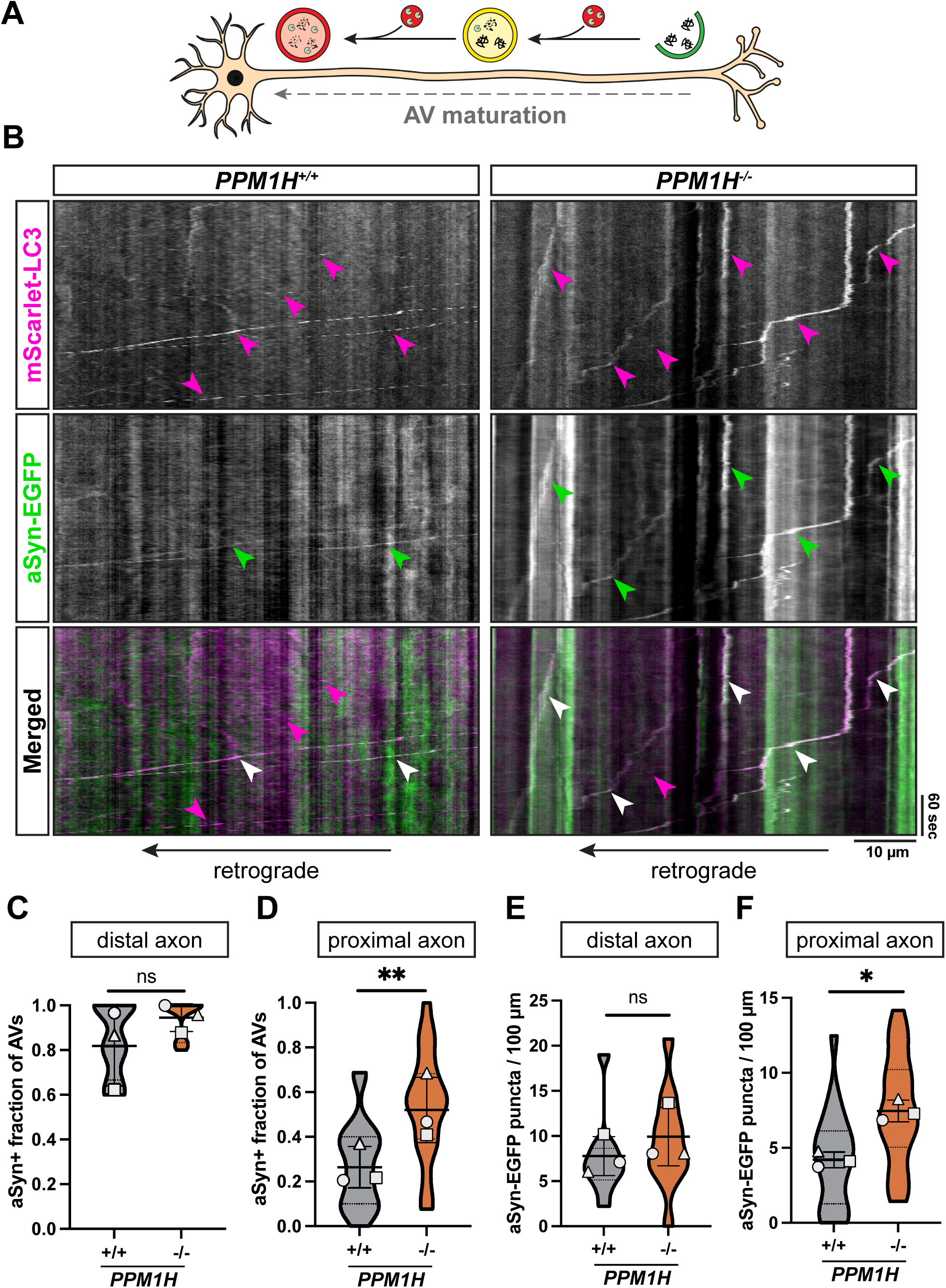
Knockout of PPM1H impairs the autophagosomal degradation of axonal aSyn-EGFP. (**A**) Schematic depicting *en route* maturation of AVs during retrograde axonal transport in WT neurons (**B**) Kymographs of mScarlet-LC3+ vesicles and aSyn-EGFP in the proximal axon of *PPM1H^+/+^* and *PPM1H^-/-^* mouse cortical neurons. Magenta arrowheads, mScarlet-LC3-positive traces; green arrowheads, aSyn- EGFP-positive traces; white arrowheads, mScarlet-LC3- and aSyn-EGFP-positive traces. (**C**) Fraction of AVs positive for aSyn-EGFP in the distal axon of *PPM1H^+/+^*and *PPM1H^-/-^* neurons (mean ± SD; n = 11 neurons from 3 independent experiments; ns, not significant, p=0.7314; mixed effects model analysis). (**D**) Fraction of AVs positive for aSyn-EGFP in the proximal axon of *PPM1H^+/+^* and *PPM1H^-/-^* neurons (mean ± SD; n = 21-22 neurons from 3 independent experiments; ** p=0.0023; mixed effects model analysis). (**E**) Number of aSyn-EGFP puncta normalized to 100 µm axonal length in the distal axon of *PPM1H^+/+^* and *PPM1H^-/-^* neurons (mean ± SD; n = 11 neurons from 3 independent experiments; ns, not significant, p=0.3484; mixed effects model analysis). (**F**) Number of aSyn-EGFP puncta normalized to 100 µm axonal length in the proximal axon of *PPM1H^+/+^*and *PPM1H^-/-^* neurons (mean ± SD; n = 21-22 neurons from 3 independent experiments; * p=0.0205; mixed effects model analysis). For panels C-F, scatter plot points indicate the means of three independent experiments.

We asked whether disruption of AV transport by *PPM1H* KO affects the autophagosomal degradation of axonal aSyn. To test this, we overexpressed aSyn-EGFP in *PPM1H^+/+^* and *PPM1H^-/-^* mouse cortical neurons together with mScarlet-LC3B as an AV marker (Figure 2B). We saw no difference in AV density in the distal axon, suggesting that de novo AV formation is not dependent on PPM1H (Figure S1E). We also noted that approximately 80-90% of AVs colocalized with aSyn-EGFP in the distal axon of both *PPM1H^+/+^* and *PPM1H^-/-^* neurons, indicating that cargo uptake into newly formed AVs was not affected by PPM1H deficiency (Figure 2C).

In the proximal axon, we observed a much lower fraction of aSyn+ AVs (Figure 2D), consistent with the degradation of engulfed aSyn-EGFP during the retrograde transport of maturing autophagosomes (Figure 2A). Notably, the fraction of aSyn+ AVs in the proximal axon of *PPM1H^-/-^* neurons was significantly higher than in *PPM1H^+/+^* neurons (Figure 2D). This was accompanied by a higher overall density of aSyn-EGFP puncta in the proximal axon, while we observed no difference in the density of aSyn-EGFP puncta in the distal axon (Figure 2E-F). In sum, these data indicate that the disruption of AV transport in neurons lacking PPM1H impairs the effective autophagosomal degradation of overexpressed aSyn, leading to its accumulation in the proximal axon.

### PFF-induced aSyn aggregation is aggravated in primary PPM1H KO neurons but not in human *PPM1H* KO or *LRRK2*-p.R1441H KI iNeurons

Previous work has shown exacerbated aSyn aggregate formation upon PFF treatment in neurons expressing hyperactive LRRK2 [29–32]. Notably, PFF-induced aSyn aggregates were observed primarily in axons [30,31]. We hypothesized that the exacerbated aSyn aggregation in neurons expressing hyperactive LRRK2 is caused by RAB hyperphosphorylation, resulting in defective AV transport and AV maturation. To investigate this, we tested whether *PPM1H* KO phenocopies the effect of hyperactive LRRK2 on aSyn aggregation. We first assessed aSyn aggregation in iPSC-derived *PPM1H^-/-^* iNeurons, which show increased levels of LRRK2-phosphorylated RAB proteins [12]. We exposed iNeurons to PFFs on DIV14 and fixed the cells two or three weeks after PFF treatment on DIV28 or DIV35 (Figure 3A). Aggregation of endogenously expressed aSyn was detected using phospho-serine 129 (pS129)-specific immunolabeling. Treatment with PFFs, but not PBS control, induced the formation of pS129-positive aSyn aggregates, the majority of which were located outside of the MAP2-positive somatodendritic compartment (Figure S2). As established in the published literature, pS129-positive area was normalized to MAP2 area as an indicator of neuronal density for quantification of aSyn aggregation levels [32,36–39]. Comparing PFF-induced aSyn aggregation in *PPM1H^-/-^* and WT iNeurons, we found no difference in pS129-positive aggregate area at both two weeks (DIV28) and three weeks (DIV35) after PFF treatment (Figure 3B-C and Figure S3). However, expression of hyperactive *LRRK2*-p.R1441H also did not result in a significant increase in pS129-positive aggregates at either timepoint (Figure 3B-C and Figure S3). Thus, neither loss of PPM1H nor expression of hyperactive LRRK2 resulted in a detectable effect in iNeurons in this assay, suggesting a limited sensitivity of this model system to detect differences in PFF-induced aSyn aggregation in the context of RAB hyperphosphorylation under the conditions used in our study.

**Figure 3.**
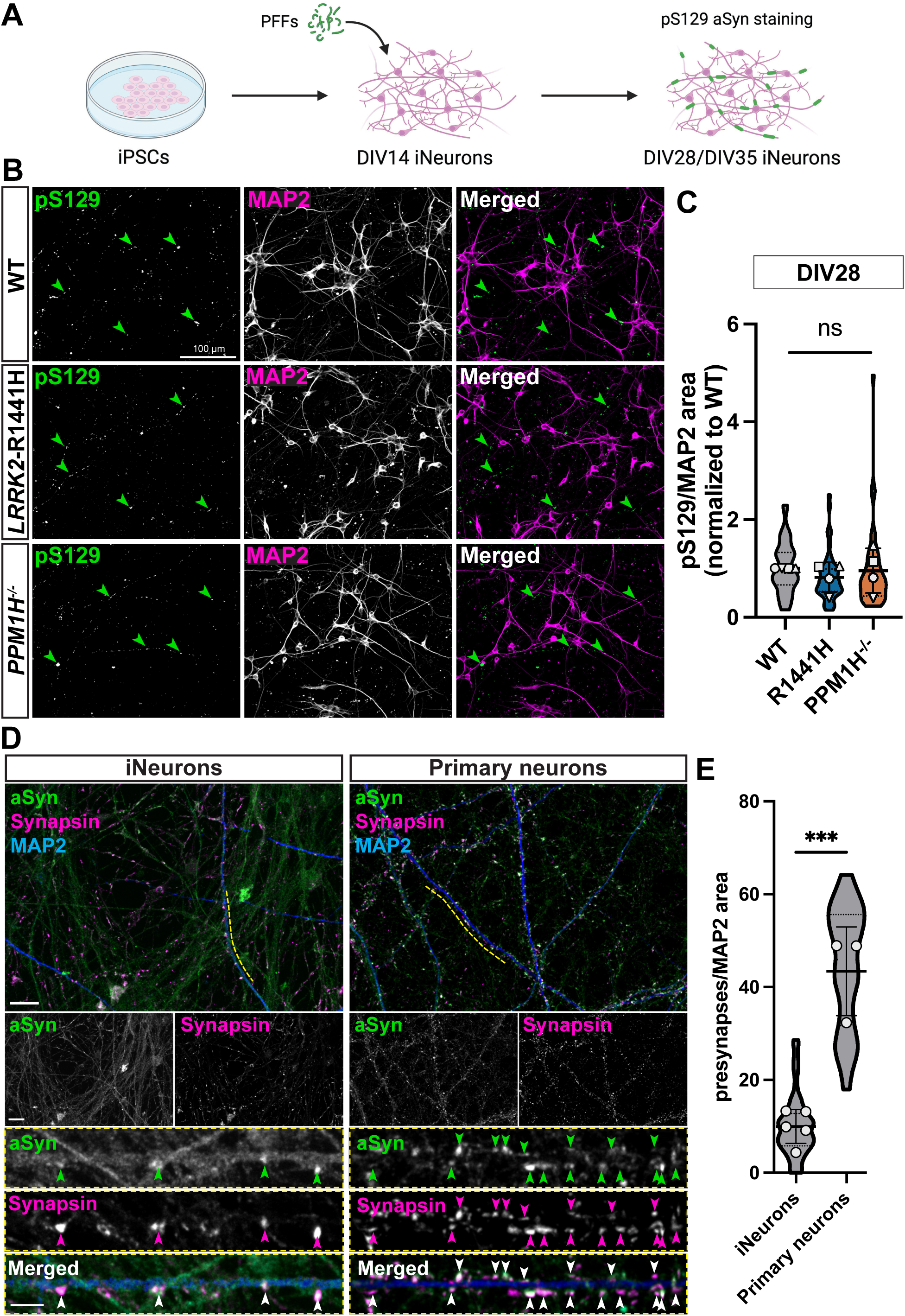
Knockout of PPM1H or knockin of *LRRK2*-p.R1441H do not alter PFF-induced aSyn aggregation in iNeurons. (**A**) Schematic depicting the experimental timeline. WT, *LRRK2*-p.R1441H KI, and *PPM1H^-/-^* iNeurons were exposed to 0.5 µM or 0.25 µM human PFFs on DIV14. iNeurons treated with 0.5 µM PFFs were fixed on DIV28 and iNeurons treated with 0.25 µM PFFs were fixed on DIV35. Fixed neurons were stained for pS129 aSyn and MAP2. (**B**) Representative images of DIV28 WT, *LRRK2*-p.R1441H KI, and *PPM1H^-/-^* iNeurons stained for pS129-positive aSyn aggregates and MAP2 after 14 days of incubation with PFFs. Green arrowheads highlight examples of pS129-positive aSyn aggregates. (**C**) pS129/MAP2 area normalized to WT control in DIV28 WT, *LRRK2*-p.R1441H KI, and *PPM1H^-/-^* iNeurons (mean ± SD; n = 44-47 fields of view from 4 independent experiments; ns, not significant, p>0.3275; mixed effects model analysis). (**D**) Representative images of DIV28 WT iNeurons and DIV21 WT mouse cortical neurons stained for aSyn (green), synapsin (magenta), and MAP2 (blue). Dashed lines indicate areas that are shown at a higher magnification in insets below. Arrowheads highlight examples of synapsin-positive presynaptic sites with enrichment of aSyn. Scale bars, 10 µm in images with larger field of view and 3 µm in insets. (**E**) Number of presynapses (defined as synapsin puncta associated with MAP2-positive dendrites) normalized to 100 µm^2^ of somatodendritic MAP2-positive area (mean ± SD; n = 3-4 independent experiments with 60-101 fields of view; *** p=0.0003; two-tailed t-test). For panels C and E, scatter plot points indicate the means of independent experiments.

One possible explanation for the lack of a response in iNeurons is that, similar to other iPSC-derived neurons, these cells exhibit slow synaptogenesis leading to a low density of bone fide synapses under conditions comparable to those in our study [40]. Consistent with this, we noted a substantially lower density of presynaptic puncta (synapsin puncta associated with MAP2-positive dendrites) in iNeurons at DIV28 as compared to primary mouse cortical neurons at DIV21 (Figure 3D-E). Of note, we further observed striking differences in the intracellular distribution of aSyn. While aSyn staining in primary neurons revealed strong enrichment of aSyn at presynaptic puncta, iNeurons displayed a more diffuse axonal aSyn signal, with only a small number of aSyn-enriched puncta colocalizing with presynapses (Figure 3D). This distribution pattern is reminiscent of the previously described localization of aSyn in immature (DIV6) primary neurons, where aSyn was primarily detected in the cytosol, with limited synaptic presence [41].

Given prior reports indicating that aSyn aggregation is dependent on synaptic activity [33], we hypothesized that the more synaptically mature primary neurons might be a more sensitive system for detecting effects of RAB hyperphosphorylation an aSyn aggregation. Primary WT and PPM*1H* KO mouse cortical neurons were exposed to PFFs on DIV7 and fixed and stained for pS129-positive aSyn aggregates on DIV21 (Figure 4A). Formation of pS129-positive aggregates was observed upon treatment with PFFs but not PBS control (Figure S4A). The majority of pS129-positive structures were located outside of MAP2-positive dendrites (Figure S4A).

**Figure 4.**
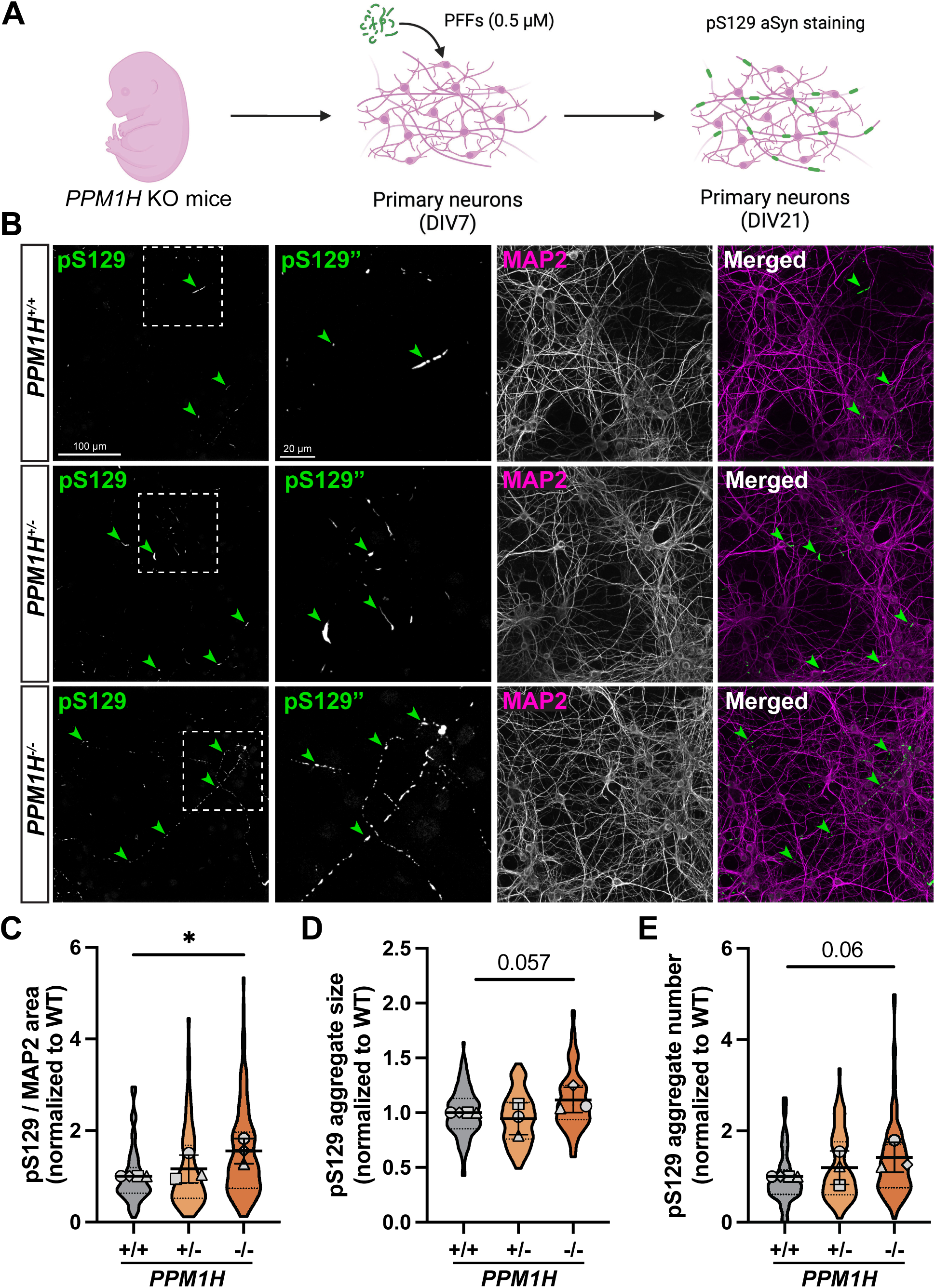
Knockout of PPM1H aggravates PFF-induced aSyn aggregation in primary neurons. (**A**) Schematic depicting the experimental timeline. *PPM1H^+/+^*, *PPM1H^+/-^*, and *PPM1H^-/-^* mouse cortical neurons were exposed to 0.5 µM human PFFs on DIV7. On DIV21, neurons were fixed and stained for pS129 aSyn and MAP2. (**B**) Representative images of DIV21 *PPM1H^+/+^*, *PPM1H^+/-^*, and *PPM1H^-/-^* neurons stained for pS129-positive aSyn aggregates and MAP2 after 14 days of incubation with PFFs. Green arrowheads highlight examples of pS129-positive aSyn aggregates. Dashed boxes indicate areas of the pS129 images that are shown at higher magnification in the second column. (**C**) pS129/MAP2 area normalized to WT control in *PPM1H^+/+^*, *PPM1H^+/-^*, and *PPM1H^-/-^* neurons (mean ± SD; n = 67-120 fields of view from 4 independent experiments; ** p=0.0125; mixed effects model analysis). (**D-E**) pS129-positive aSyn aggregate size (D) and pS129-positive aSyn aggregate number (E) normalized to WT control in *PPM1H^+/+^*, *PPM1H^+/-^*, and *PPM1H^-/-^* neurons (mean ± SD; n = 67-120 fields of view from 4 independent experiments; p values indicated in figure; mixed effects model analysis). For panels C-E, scatter plot points indicate the means of each independent experiment.

Strikingly, quantification of pS129-positive aggregates revealed an increase in pS129-positive area in *PPM1H^-/-^* neurons as compared to *PPM1H^+/+^* controls (Figure 4B-C). Furthermore, *PPM1H^+/-^*neurons showed a non-significant trend toward towards higher aggregate levels (Figure 4B-C). There was no difference in the area of MAP2-positive neurites between the different genotypes (Figure S4B-C), indicating that *PPM1H* KO did not affect neuron viability upon PFF exposure. Quantification of the size and number of individual pS129-positive aggregates showed strong trends towards bigger and more frequent aggregates in *PPM1H^-/-^* neurons, suggesting that the significant increase in overall pS129-positive area resulted from an increase in both aggregate size and number (Figure 4D-E). Together, our data show that primary *PPM1H^-/-^* neurons phenocopy the aggravated aSyn aggregation previously reported in neurons expressing hyperactive LRRK2 [29–32].

### PPM1H KO does not affect intracellular PFF uptake in primary neurons

One possible explanation for the increased formation of pS129-positive aggregates in *PPM1H^-/-^* primary neurons is increased intracellular uptake of PFFs. To test this, we treated *PPM1H^+/+^* and *PPM1H^-/-^*primary neurons with ATTO 488-labeled PFFs and measured the internalization of fluorescently labeled PFFs. Neurons were exposed to ATTO 488-labeled PFFs on DIV7 and fixed on DIV8 after washing with diluted trypsin to remove extracellular PFFs as previously described (Figure 5A) [42]. We did not observe a difference in the integrated density of internalized PFFs in the soma between *PPM1H^+/+^* and *PPM1H^-/-^* neurons (Figure 5B-C). Similarly, we found no difference in the area (Figure S5A) or fluorescence intensity of internalized PFFs (Figure S5B). Thus, the increased levels of pS129-positive aSyn aggregates in *PPM1H^-/-^*primary neurons are not caused by an enhanced intracellular uptake of PFFs. This is consistent with previous results showing no effect of the expression of hyperactive LRRK2 on neuronal PFF internalization [31], and together with our other data supports a model in which exacerbated PFF-induced aSyn aggregation may instead be caused by impaired axonal autophagy and defective aSyn degradation.

**Figure 5.**
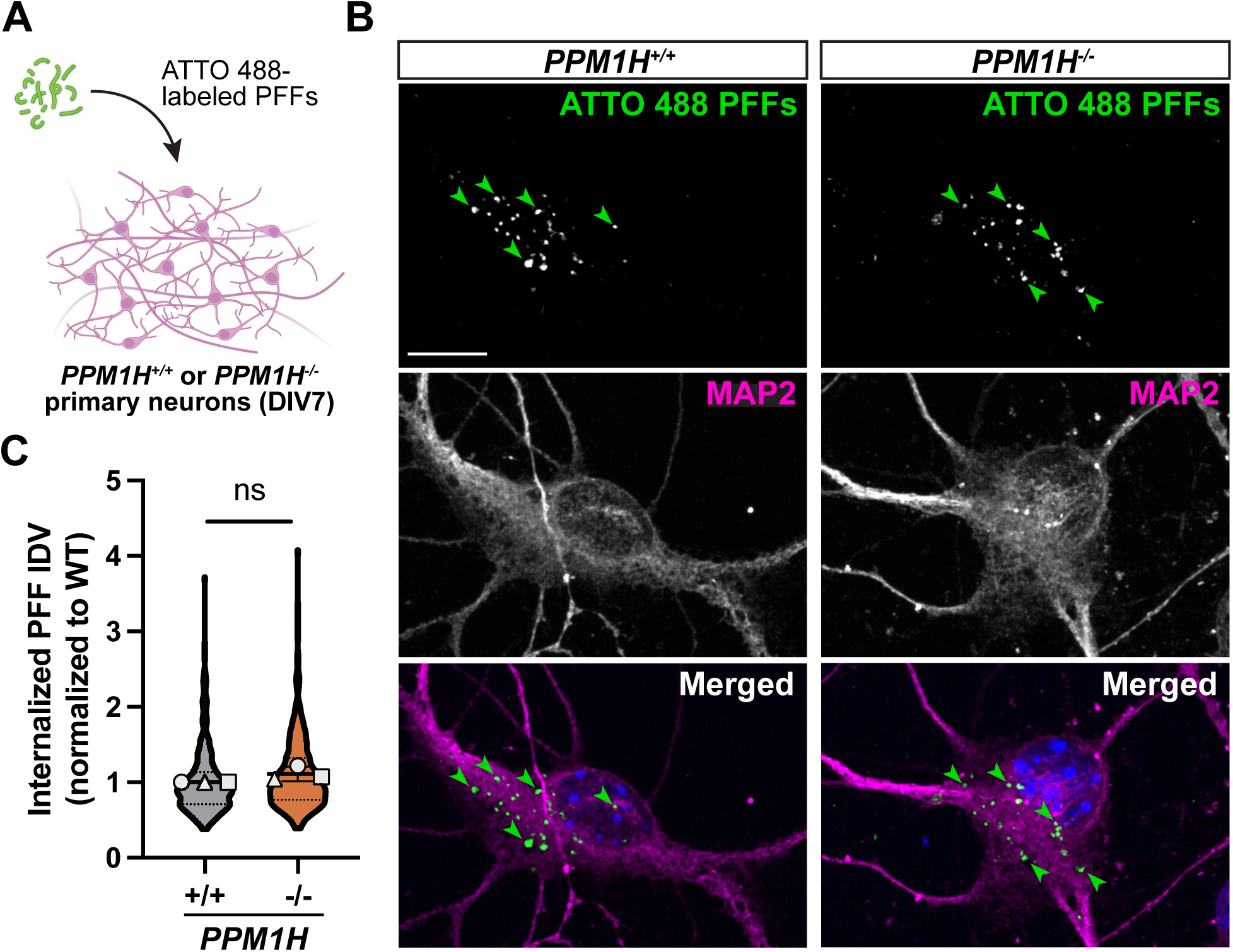
Knockout of PPM1H does not alter intracellular uptake of fluorescently labeled PFFs. (**A**) Schematic depicting the experimental timeline. *PPM1H^+/+^* and *PPM1H^-/-^*mouse cortical neurons were exposed to 0.5 µM ATTO 488-labeled PFFs on DIV7, then fixed on DIV8 and stained for MAP2. (**B**) Representative images of DIV8 *PPM1H^+/+^* and *PPM1H^-/-^* mouse cortical neurons exposed to ATTO 488- labeles PFFs for 24 h and stained for MAP2. Green arrowheads highlight examples of internalized PFFs. Scale bar, 10 µm. (**C**) Integrated density of internalized ATTO 488-labeled PFFs normalized to WT control in *PPM1H^+/+^* and *PPM1H^-/-^*neurons (n = 270 – 301 neurons from 3 independent experiments; ns, not significant, p=0.1771; mixed effects model analysis). For panel C, scatter plot points indicate the means of three independent experiments.

### PFF-induced aSyn aggregation in primary PPM1H KO neurons is reduced by LRRK2 kinase inhibition

To investigate whether the effect of *PPM1H* KO on aSyn aggregation is mediated through a LRRK2-dependent pathway, we next applied MLi-2 to *PPM1H^-/-^* primary neurons to pharmacologically inhibit LRRK2 kinase activity. *PPM1H^-/-^* neurons were treated with MLi-2 or DMSO control on DIV5, exposed to PFFs on DIV7, and fixed and stained for pS129 aggregates on DIV21 (Figure 6A). Western blots confirmed reduction of pT73 Rab10 levels in *PPM1H^-/-^* neurons treated with MLi-2 from DIV5 to DIV21 (Figure 6B). Quantification of pS129-positive area revealed that pharmacological LRRK2 kinase inhibition with MLi-2 significantly reduced the levels of pS129-positive aggregates in *PPM1H^-/-^* neurons after PFF treatment compared to DMSO control (Figure 6C-D). Treatment with MLi-2 did not affect MAP2 area in *PPM1H^-/-^* neurons (Figure S5C). These results demonstrate that loss of PPM1H increases aSyn aggregation in primary neurons in a LRRK2 kinase activity-dependent manner.

**Figure 6.**
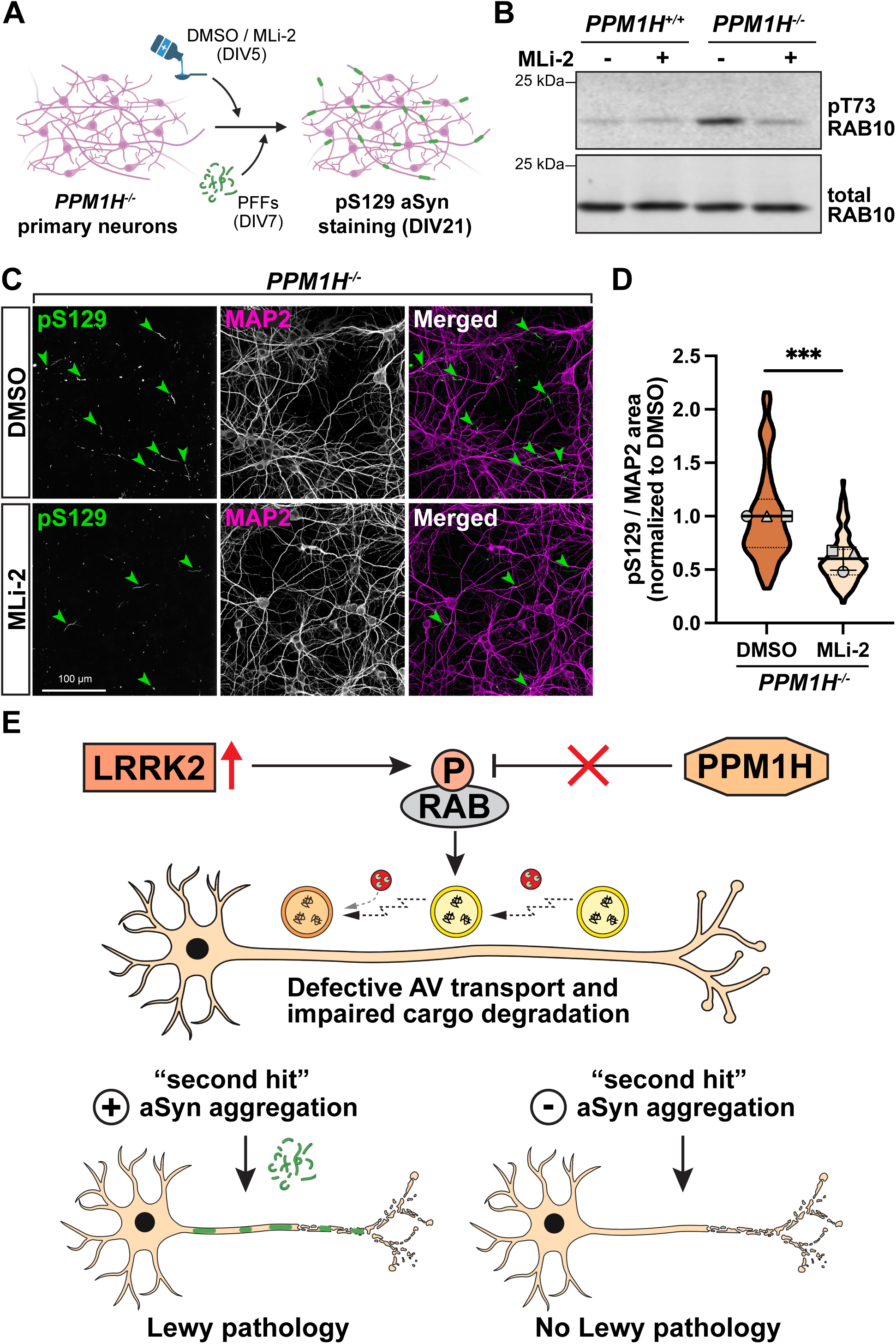
LRRK2 kinase inhibition ameliorates PFF-induced aSyn aggregation in primary PPM1H knockout neurons. (**A**) Schematic depicting the experimental timeline. *PPM1H^-/-^*mouse cortical neurons were treated with DMSO control or 300 nM MLi-2 on DIV5 and fed with media containing DMSO or MLi-2 until fixation. On DIV7, neurons were exposed to 0.5 µM human PFFs. On DIV21, neurons were fixed and stained for pS129 aSyn and MAP2. (**B**) Western blot of pT73 Rab10 and total Rab10 in DIV21 *PPM1H^+/+^* and *PPM1H^-/-^* mouse cortical neurons. Neurons were treated with DMSO or 300 nM MLi-2 on DIV5, then fed with media containing DMSO or MLi-2 until cell lysis on DIV21. (**C**) Representative images of DIV21 *PPM1H^-/-^*neurons treated with DMSO or MLi-2 and stained for pS129-positive aSyn aggregates and MAP2 after 14 days of incubation with PFFs. Green arrowheads highlight examples of pS129-positive aSyn aggregates. (**D**) pS129/MAP2 area normalized to WT control in *PPM1H^-/-^* neurons treated with DMSO or MLi-2 (mean ± SD; n = 98 fields of view from 3 independent experiments; *** p<0.001; mixed effects model analysis). Scatter plot points indicate the means of three independent experiments. (**E**) Model depicting the effect of RAB hyperphosphorylation on axonal autophagy and aSyn pathology. RAB hyperphosphorylation, resulting from either expression of hyperactive LRRK2 or loss of PPM1H, causes impaired axonal AV transport and defective degradation of autophagosomal cargo such as aSyn. Additional internal or external risk factors that promote aSyn aggregation lead to the development of Lewy pathology, while other risk factors trigger a different disease pathway, resulting in neurodegeneration without Lewy pathology.

## DISCUSSION

Inappropriate aSyn aggregation has long been considered a fundamental feature of PD pathogenesis, with accumulating evidence implicating pathogenic LRRK2 hyperactivity. However, the underlying mechanisms that may link LRRK2 hyperactivity to aSyn pathology have not been fully elucidated. Here, we show in mouse primary neurons that knockout of the LRRK2-counteracting RAB phosphatase PPM1H disrupts axonal AV transport, impairs the autophagosomal degradation of overexpressed aSyn, and exacerbates aSyn aggregate formation upon exposure to PFFs in a LRRK2 kinase-dependent manner. Our results demonstrate that loss of PPM1H phenocopies the impairment of axonal autophagy and increase in aSyn aggregation that is induced by hyperactive LRRK2, strongly indicating that both phenotypes are mediated by hyperphosphorylation of RAB GTPases [11,12,29–31].

We found that axonal AV transport was markedly impaired in primary neurons from *PPM1H* KO mice. Treatment with the selective LRRK2 inhibitor MLi-2 rescued AV motility, confirming that the disruption of AV transport induced by loss of PPM1H is dependent on LRRK2 kinase activity. Homozygous primary *PPM1H^-/-^* neurons showed a more pronounced disruption of AV transport compared to heterozygous *PPM1H^+/-^* neurons, with a higher fraction of AVs remaining fully stationary during time-lapse acquisition and a longer overall fraction of time that AVs spent paused. This is consistent with the observation that the strongly hyperactivating *LRRK2*-p.R1441H mutation disrupts AV transport to a greater extent than the moderately hyperactivating *LRRK2*-p.G2019S mutation [12], and further establishes that the magnitude of AV transport deficits scales with the level of LRRK2-mediated RAB hyperphosphorylation. Notably, we found that the more pronounced disruption of AV transport in *PPM1H^-/-^* primary neurons correlated with a stronger effect on aSyn aggregation compared to *PPM1H^+/-^* neurons.

Several previous studies have shown that defective retrograde AV transport impairs AV maturation and fusion with lysosomal vesicles, disrupting effective local degradation of autophagosomal cargo. Defective AV transport caused by knockin of the *LRRK2*-p.G2019S mutation or knockdown of the motor adaptor proteins Huntingtin or JIP1 was associated with impaired acidification or impaired degradation of the autophagosomal cargos of axonal AVs [11,19,21]. To investigate whether impaired AV transport affects the autophagosomal degradation of aSyn, we challenged primary *PPM1H* KO neurons by overexpressing aSyn. We found that defective AV transport in *PPM1H^-/-^* neurons was accompanied by both an increased fraction of AVs containing aSyn-EGFP and an increased density of aSyn-EGFP puncta in the proximal axon. Since we observed no difference between WT and *PPM1H^-/-^* neurons in the distal axon, these results strongly indicate that loss of PPM1H does not affect autophagosome biogenesis or cargo uptake, but does impair the autophagosomal degradation of aSyn during axonal transport. Consistent with our findings, a previous report found that inhibition of AV maturation by disrupting the axonal delivery of lysosomal vesicles induced aSyn accumulation in the axon [18].

The expression of hyperactive LRRK2 has been shown to exacerbate the formation of pS129-positive aSyn aggregates upon PFF treatment in a number of previous studies [29–32]. We found that primary *PPM1H^-/-^*neurons phenocopied this effect of pathogenic LRRK2, exhibiting increased pS129-positive aSyn aggregation after exposure to PFFs as compared to WT control. In contrast, neither the loss of PPM1H nor the expression of hyperactive LRRK2 resulted in a detectable effect on aSyn aggregation in iNeurons. This interesting discrepancy may be explained by notable differences between iNeurons and primary neurons. First, we found that compared to abundant synapses and strong enrichment of aSyn at presynapses in primary neurons, iNeurons exhibited fewer synapses and a more diffuse intraaxonal distribution of aSyn. It is possible that aSyn enrichment at the presynapse, a major site of autophagosome biogenesis, increases the burden on aSyn degradation via axonal autophagy, making primary neurons more susceptible to defects in this pathway. Consistent with this interpretation, previous work has found that increased synaptic activity exacerbates the detrimental effect of hyperactive LRRK2 on aSyn aggregation [33]. Second, we observed higher levels of axonal branching in primary cortical neurons compared to iNeurons in this and previous work [43]. Extensive axonal branching results in a longer overall length of the axonal arbor, which is predicted to amplify the detrimental effects of defective AV transport and may thus explain a higher vulnerability of primary neurons to the transport deficits resulting from RAB hyperphosphorylation. Together, our results indicate that iNeurons, while remaining a highly valuable model for observing and characterizing changes in axonal transport dynamics, have limited sensitivity in detecting downstream effects of RAB hyperphosphorylation on aSyn aggregation under the culture conditions used here.

Mechanistically, we found that aSyn aggregation in primary *PPM1H^-/-^*neurons was attenuated by pharmacological LRRK2 kinase inhibition, confirming that the exacerbated aSyn aggregation induced by loss of PPM1H is dependent on LRRK2-mediated RAB hyperphosphorylation. Furthermore, we observed no difference in the intracellular uptake of fluorescently labeled PFFs between primary *PPM1H^-/-^* neurons and WT neurons. Similarly, previous work found no effect of hyperactive LRRK2 on neuronal PFF internalization [31], suggesting that increased aSyn aggregation in the context of LRRK2-mediated RAB hyperphosphorylation is not caused by an enhanced intracellular uptake of PFFs. Instead, our data support a model in which defective axonal AV transport and impaired axonal aSyn degradation caused by RAB hyperphosphorylation result in an increased neuronal vulnerability to PFF-induced aSyn aggregation. This is supported by previous studies demonstrating that PFF-induced aSyn aggregation primarily occurs in the axon [30,31]. We cannot fully exclude the potential contribution of lysosomal defects, although we observed no change in density, transport, or acidification of axonal LAMP1 vesicles in *LRRK2*-p.G2019S KI neurons in previous work [11].

An interesting neuropathological finding is that about 20-50% of LRRK2-PD cases do not show aSyn aggregation to Lewy bodies and Lewy neurites [44]. One possible explanation is that LRRK2 mutations and the resulting intracellular defects predispose carriers to multiple disease pathways rather than a single one. Certain internal or external risk factors may promote aSyn aggregation and lead to Lewy pathology, while others may trigger a different disease pathway, resulting in neurodegeneration without Lewy pathology (Figure 6E) [45]. Our data presented here fit to this model, as both aSyn overexpression and PFF exposure represent a “second hit” that challenges axonal degradation in neurons with elevated levels of LRRK2-phosphorylated RAB proteins. In the absence of an additional stressor, impaired baseline degradation of aSyn may be compensated by increased aSyn secretion [22] and subsequent extracellular degradation. Instead, a different “second hit” may cause neurodegeneration through another aSyn-independent disease pathway (Figure 6E). Notably, this model is consistent with the known incomplete penetrance of LRRK2 mutations [46].

An increasing number of studies have linked LRRK2 hyperactivity to the neurodegeneration of PD, leading to the development of pharmacological LRRK2 kinase inhibitors. In this study, we show in primary neurons that loss of the LRRK2-counteracting RAB phosphatase PPM1H disrupts the axonal transport of autophagosomes, impairs the autophagosomal degradation of axonal aSyn, and exacerbates the PFF-induced aggregation of aSyn in a LRRK2 kinase-dependent manner. Our results strongly suggest that the previously reported effects of hyperactive LRRK2 on autophagy and aSyn pathology are mediated by hyperphosphorylation of RAB GTPases. Unraveling the underlying mechanism of how LRRK2 hyperactivity causes neurodegeneration has important translational implications, as side effects reported for LRRK2 kinase inhibitors may necessitate more specific therapeutic approaches.

## MATERIALS AND METHODS

**Table 1.** Key resources.

| REAGENT or RESOURCE | SOURCE | IDENTIFIER |
| --- | --- | --- |
| **Antibodies** |  |  |
| Anti-RAB10 antibody, Mouse Monoclonal | Nanotools | 0680-100/Rab10-605B11, RRID: AB_2921226 |
| Anti-RAB10 (phospho T73) antibody, Rabbit Monoclonal | Abcam | ab230261, RRID: AB_2811274 |
| Anti-PPM1H antibody, Sheep Polyclonal | MRC PPU reagents and services, University of Dundee, Scotland | DA018, 3<sup>rd</sup> bleed, RRID: AB_2923281 |
| Anti-Rabbit IgG-IRDye 800CW, Donkey Polyclonal | Li-COR Biosciences | 926-32213, RRID: AB_621848 |
| Anti-Mouse IgG-IRDye 680RD, Donkey Polyclonal | Li-COR Biosciences | 926-68073, RRID: AB_10954442 |
| Anti-Sheep Alexa Fluor 680, Donkey Polyclonal | Thermo Fisher | A21102, RRID: AB_10374469 |
| Anti-αSynuclein (phospho S129) antibody, Rabbit Monoclonal | Abcam | ab51253, RRID: AB_869973 |
| Anti-αSynuclein antibody, Mouse Monoclonal | BD | 610787, RRID: AB_398108 |
| Anti-Synapsin1/2 antibody, Rabbit Polyclonal | Synaptic Systems | 106002, RRID: AB_887804 |
| Anti-MAP2 antibody, Mouse Monoclonal | Millipore | MAB3418, RRID: AB_94856 |
| Anti-MAP2 antibody, Chicken Polyclonal | Abcam | ab5392, RRID: AB_2138153 |
| Alexa Fluor secondary antibody for pS129 (IF)<br>Anti-Rabbit Alexa Fluor 488, Goat Polyclonal | Thermo Fisher | A-11034, RRID: AB_2576217 |
| Alexa Fluor secondary antibody for MAP2 (IF)<br>Anti-Mouse Alexa Fluor 594, Goat Polyclonal | Thermo Fisher | A-11005, RRID: AB_2534073 |
| **Chemicals, Peptides, and Recombinant Proteins** |  |  |
| MLi-2 | Tocris | 5756 |
| DMSO | AppliChem | A3672,0100 |
| Poly-L-Lysine (mol wt 70,000 – 150,000) | Sigma-Aldrich | P1274 |
| HBSS (10×) | Thermo Fisher | 14185-052 |
| 1M HEPES | Thermo Fisher | 15630-080 |
| 2.5% Trypsin | Thermo Fisher | 15090-046 |
| Minimum essential medium (MEM) | Thermo Fisher | 11095-072 |
| Horse serum (heat inactivated) | Thermo Fisher | 16050-122 |
| Sodium pyruvate | Thermo Fisher | 11360-039 |
| D-Glucose solution 45% | Sigma-Aldrich | G8769 |
| GlutaMAX | Thermo Fisher | 35050061 |
| B27 Supplement | Thermo Fisher | 17504-044 |
| Neurobasal Medium | Thermo Fisher | 21103-049 |
| Penicillin-Streptomycin | Thermo Fisher | 15140-122 |
| Lipofectamine 2000 Transfection Reagent | Thermo Fisher | 11668019 |
| Matrigel Growth Factor Reduced | Corning | 354230 |
| Essential 8 Medium | Thermo Fisher | A1517001 |
| ReLeSR | Stemcell Technologies | 05872 |
| Accutase | Sigma-Aldrich | A6964 |
| ROCK Inhibitor Y-27632 | Selleckchem | S1049 |
| Knockout Serum Replacement | Thermo Fisher | 10828010 |
| DMEM/F-12, HEPES | Thermo Fisher | 11330032 |
| N2 Supplement | Thermo Fisher | 17502048 |
| Non-essential Amino Acids (NEAA) | Thermo Fisher | 11140050 |
| Doxycycline | Sigma-Aldrich | D9891 |
| Poly-L-Ornithine | Sigma-Aldrich | P3655 |
| BrainPhys Neuronal Medium | Stemcell Technologies | 05790 |
| Laminin | Corning | 354232 |
| BDNF | PeproTech | 450-02 |
| NT-3 | PeproTech | 450-03 |
| Microcystin-LR | Sigma-Aldrich | 475815 |
| Halt Protease and Phosphatase Inhibitor Cocktail | Thermo Fisher | 78442 |
| 5-Fluoro-2'-deoxyuridine | Sigma-Aldrich | F0503 |
| Uridine | Sigma-Aldrich | U3003 |
| EveryBlot Blocking Buffer | BioRad | 12010020 |
| **Critical Commercial Assays** |  |  |
| BCA Protein Assay Kit | Thermo Fisher | 23225 |
| ZymoPURE II Plasmid Maxiprep Kit | ZymoResearch | D4203 |
| **Experimental Models: Cell Lines** |  |  |
| Human: KOLF2.1J WT iPSCs | B. Skarnes (Jackson Laboratories, Connecticut) | RRID: CVCL_B5P3 |
| Human: KOLF2.1J LRRK2-R1441H iPSCs | B. Skarnes (Jackson Laboratories, Connecticut) | N/A |
| Human: KOLF2.1J PPM1H KO iPSCs | Dou et al., 2023 [12] | RRID: CVCL_C7TY |
| **Experimental Models: Organisms/Strains** |  |  |
| Mouse: LRRK2-G2019S knock-in | Taconic | 13940 |
| Mouse: C57BL/6NTac WT | Taconic | B6 |
| Mouse: PPM1H knock-out | Taconic | TF3142 |
| **Recombinant DNA** |  |  |
| Plasmid: CMV mScarlet-LC3B | Modified from EGFP-LC3B (gift from T. Yoshimori, Osaka University, Japan) | Addgene plasmid #200431 |
| Plasmid: CMV aSyn-EGFP | VectorBuilder, same linker as described in [47] | VB230227-1242vtx |
| Plasmid: PB-TO-hNGN2 | Gift from iPSC Neurodegenerative Disease Initiative (iNDI) & Michael Ward | Addgene plasmid #172115 |
| Plasmid: piggyBac™ transposase vector | Transposagen | N/A |
| **Software and Algorithms** |  |  |
| FIJI (Release 2.9.0) | NIH, USA | http://fiji.sc |
| Prism 10 | GraphPad | https://www.graphpad.com/scientific-software/prism/ |
| RStudio: Integrated Development for R (2022.07.0) | RStudio Team | http://www.rstudio.com/ |
| R package: nlme | Pinheiro J, Bates D, R Core Team | http://CRAN.R-project.org/package=nlme |
| R package: lme4 | Bates D, Mächler M, Bolker B, Walker S | https://cran.r-project.org/web/packages/lme4/index.html |
| Matlab R2022a | MathWorks | https://www.mathworks.com/products/matlab.html |
| KymoSuite (custom Matlab script) | Guedes-Dias et al., 2019 [48] | https://github.com/jnirschl/kinesin-3_guedes-dias_2018/tree/master/kymoSuite,<br>https://zenodo.org/record/2530934 |
| VisiView | Visitron | https://www.visitron.de/products/visiviewr-software.html |
| LI-COR Image Studio | LI-COR | https://www.licor.com/bio/image-studio/ |
| Adobe Illustrator 2023 | Adobe | https://www.adobe.com/products/illustrator.html |
| BioRender | BioRender | https://biorender.com/ |
| **Other** |  |  |
| 35 mm #1.5 glass bottom imaging dishes | MatTek | P35G-1.5-20-C |
| 12 mm #1.5 glass coverslips | R. Langenbrinck GmbH | 01-0012/5 |
| ProLong Gold Antifade Mountant | Thermo Fisher | P36930 |

### Primary neuron culture

*PPM1H* KO mice (Taconic #TF3142) were provided by Dario Alessi, University of Dundee, and have been previously described [34]. A license for maintaining a colony of PPM1H KO mice was obtained from the manufacturer. All animal experiments were approved by the local animal research council and complied with the legislation for the management and care of laboratory animals in the state of Lower Saxony, Germany. Mouse cortices were dissected from littermate PPM1H^+/+^, PPM1H^+/-^, and PPM1H^-/-^ embryos of either sex at day 15.5. Upon sacrificing PPM1H KO embryos, tails were collected and used for genotyping. Cortical neurons were isolated by digestion with 0.25% trypsin and trituration through a small-bore serological pipette. Neurons were plated in attachment media (MEM supplemented with 10% horse serum, 33 mM D-glucose and 1 mM sodium pyruvate) on 35 mm glass-bottom imaging dishes (P35G-1.5-20-C; MatTek) for live-imaging experiments (200,000 neurons per dish), 24-well plates with 12 mm #1.5 coverslips for PFF experiments (100,000 neurons per well), or 6-well plates for biochemistry experiments (1,000,000 neurons per well). Attachment media was replaced with maintenance media (Neurobasal supplemented with 2% B-27, 33 mM D-glucose, 2 mM GlutaMAX, 100 U/mL penicillin and 100 mg/mL streptomycin) 3-5 hours after plating. In imaging dishes and 6-well plates, 40% of the media was replaced with fresh maintenance media twice per week. Neurons in 24-well plates used for PFF experiments were fed by adding 10% maintenance media twice per week.

### PiggyBac-mediated iPSC-derived neuron differentiation and iNeuron culture

KOLF2.1J-background WT, LRRK2-p.R1441H KI, and PPM1H KO iPSCs have been described and characterized previously [12]. iPSCs were cultured on plates coated with Growth Factor Reduced Matrigel (Corning) and fed daily with Essential 8 media. To stably express doxycycline-inducible hNGN2 using a PiggyBac delivery system, iPSCs were transfected with PB-TO-hNGN2 vector (gift from M. Ward, NIH, Maryland) in a 1:2 ratio (transposase:vector) using Lipofectamine Stem [12]. After 72 hours, transfected iPSCs were selected for 48 hours with 0.5 mg/mL puromycin. Differentiation of PB-TO-hNGN2 iPSCs into iNeurons was then performed using an established protocol [12,49]. In brief, iPSCs were passaged using Accutase and plated on Matrigel-coated dishes in induction media (DMEM/F12 supplemented with 1% N2-supplement, 1% NEAA, and 1% GlutaMAX, and containing 2 mg/mL doxycycline). After 72 hours of doxycycline exposure, iNeurons were dissociated with Accutase and cryo-preserved in liquid N2.

Cryo-preserved, pre-differentiated iNeurons were thawed and plated on coverslips coated with poly-L-ornithine at a density of 100,000 neurons per well in 24-well plates. iNeurons were cultured in BrainPhys Neuronal Media (StemCell) supplemented with 2% B-27, 10 ng/mL BDNF (PeproTech), 10 ng/mL NT-3 (PeproTech), and 1 mg/mL laminin. Neurons were fed by adding 10% fresh media twice per week. 10 mM 5-Fluoro-20-deoxyuridine and 10 mM uridine were included at the time of plating to prevent survival of mitotic cells. These drugs were removed 24 hours after plating. For each experimental condition, cells from at least two different batches of induction were used over three or more independent experimental cultures. Protocols for PiggyBac-mediated differentiation of iPSCs and culture of iNeurons can be found on https://doi.org/10.17504/protocols.io.e6nvwj54dlmk/v1 and https://doi.org/10.17504/ protocols.io.x54v9dj4zg3e/v1.

### Live-cell imaging

Live-cell imaging experiments were performed at the Live-Cell Imaging Facility of the Max Planck Institute for Multidisciplinary Sciences, Goettingen. Primary mouse cortical neurons were imaged on DIV8 in low fluorescence Hibernate E medium (Brain Bits) supplemented with 2% B27 and 2 mM GlutaMAX. Time lapse series were recorded on a Visitron CSU-W1 Spinning Disk Confocal system with a Nikon Ti2 inverted microscope using a Plan Apochromat 60x 1.40 NA oil immersion objective and a Prime BSI sCMOS camera controlled by VisiView software. Axons were identified based on morphological parameters [43,50]. For AV motility experiments, time lapse recordings were acquired at a frame rate of 1 frame/sec for 5 minutes in the mid-axon (> 300 µm from the soma and > 100 µm from the distal axon terminal). For analyzing the fraction of aSyn-EGFP-positive AVs, time lapse series were recorded at a frame rate of 1 frame/sec for 5 minutes in the proximal axon (< 150 µm from the cell soma).

### PFF assay

Purification of recombinant aSyn and generation of aSyn PFFs was performed as previously described [51]. PFFs at a concentration of 25 µM were sonicated for 30 sec using a Bandelin SonoPlus sonicator (10% power, 1 cycle) immediately before usage. PFFs were then diluted in media to 2x of their final concentration and added to cultured neurons by removing 50% of culture media from each well and replacing it with an equal amount of PFF-containing media. For PBS control conditions, an equal volume of PBS was used instead of PFFs. PFFs were added to DIV7 mouse cortical neurons at a final concentration of 0.5 µM and to DIV14 iNeurons at a final concentration of 0.25 or 0.5 µM. After adding PFFs, neurons were fed by adding 10% of fresh media twice per week. For MLi-2 experiments, mouse cortical neurons were treated with DMSO or 300 nM MLi-2 on DIV5 and fed with media containing DMSO or MLi-2 until fixation. On DIV21 (mouse cortical neurons) or DIV28/DIV35 (iNeurons), cells were fixed with 4% PFA and 4% sucrose at 37°C for 9 minutes. After three washes with PBS, cells were blocked/permeabilized for 1.5 hours with PBS containing 5% normal goat serum, 1% BSA, and 0.1% Triton-X. Neurons were then incubated with primary antibodies diluted in blocking solution overnight at 4°C, washed three times with PBS, and incubated in secondary antibodies diluted in blocking solution for 1 hour at RT. The following primary antibodies were used: Abcam EP1536Y Rabbit anti-pS129 alpha-synuclein (ab51253; 1:500), Millipore Mouse anti-MAP2 (MAB3418; 1:200). After three washes with PBS and nuclear counterstaining with DAPI, coverslips were mounted in ProLong Gold Antifade mountant. Immunofluorescently stained mouse cortical neurons were imaged on the Visitron CSU-W1 Spinning Disk confocal system described above using a 40x Plan Apochromat 0.95 NA air objective. Images were recorded as z-stacks with 400 nm step size. iNeurons were imaged on a Zeiss Axioplan 2 epifluorescence microscope controlled by ZEN black software using a 20x Plan Apochromat 0.80 NA air objective and Zeiss Apotome 3.

### ATTO 488-labeled PFF assay

ATTO 488-labeled PFFs were generously provided by T. Outeiro, University Medical Center Goettingen. Labeled PFFs were diluted to 25 µM with PBS and sonicated as described above. Labeled PFFs were then diluted in media to 1 µM and added to cultured mouse cortical neurons on DIV7 by removing 50% of culture media from each well and replacing it with an equal amount of PFF-containing media (final concentration: 0.5 µM). On DIV8, neurons were washed once with 1x Trypsin-EDTA diluted 1:10 in PBS for 30 sec and once with PBS to remove extracellular PFFs, as described previously [42]. Cells were then fixed with 4% PFA and 4% sucrose and stained for MAP2 as described above. Stained neurons were imaged on the Visitron CSU-W1 Spinning Disk confocal system described above using a 100x Apochromat 1.49 NA oil immersion objective. Images were recorded as z-stacks with 300 nm step size.

### Immunofluorescence

DIV28 iNeurons and DIV21 primary cortical neurons were fixed were with 4% PFA and 4% sucrose at 37°C for 9 minutes. After three washes with PBS, cells were blocked/permeabilized for 1.5 hours with PBS containing 5% normal goat serum, 1% BSA, and 0.1% Triton-X. Neurons were then incubated with primary antibodies diluted in blocking solution overnight at 4°C, washed three times with PBS, and incubated in secondary antibodies diluted in blocking solution for 1 hour at RT. The following primary antibodies were used: Synaptic Systems Rabbit anti-synapsin (#106002; 1:500), BD Mouse anti-aSyn (#610787; 1:500), Abcam Chicken anti-MAP2 (ab5392; 1:5000). After three washes with PBS and nuclear counterstaining with DAPI, coverslips were mounted in ProLong Gold Antifade mountant. Immunofluorescently stained neurons were imaged on the Visitron CSU-W1 Spinning Disk confocal system described above using a 100x Apochromat 1.49 NA oil objective. Images were recorded as z-stacks with 300 nm step size.

### Immunoblotting

Neurons were washed twice with ice cold PBS and lysed with RIPA buffer (50 mM Tris-HCl, 150 mM NaCl, 0.1% Triton X-100, 0.5% deoxycholate, 0.1% SDS, 2x Halt Protease and Phosphatase inhibitor, 2mg/mL microcystin-LR). Samples were centrifuged for 10 min at 16,200 g, and protein concentration of the supernatant was determined by BCA assay. Proteins were resolved on 10% (PPM1H) or 15% (Rab proteins) acrylamide gels. Proteins were transferred to Immobilon-FL PVDF membranes (Millipore) using a wet blot transfer system. For PPM1H blots, membranes were then stained for total protein using LI-COR Revert 700 Total Protein Stain. Following imaging of total protein stain, membranes were de-stained, blocked for 5 minutes with Bio-Rad EveryBlot blocking buffer, and incubated with primary antibody diluted in EveryBlot for 90 minutes at room temperature. For pT73 RAB10 and total RAB10 blots, membranes were blocked for 5 minutes with EveryBlot after protein transfer and then incubated with primary antibody diluted in EveryBlot at 4°C overnight. After three washes with TBS (50 mM Tris-HCl [pH 7.4], 274 mM NaCl, 9 mM KCl) with 0.1% Tween-20, membranes were incubated with secondary antibodies diluted in EveryBlot with 0.01% SDS for 1 hr at RT. Following three more washes with TBS with 0.1% Tween-20, membranes were imaged using an Odyssey CLx Infrared Imaging System (LI-COR). Western blots were analyzed with Image Studio Software (LI-COR).

### Quantification and statistical analysis

#### AV motility

Kymographs of LC3+ AVs were generated with the Multiple Kymograph plugin for FIJI using a line width of 5 pixels. Vesicle tracks were traced manually with a custom MATLAB GUI (KymoSuite). Motile AVs were scored as anterograde (net displacement > 10 µm in the anterograde direction within 5 minutes), retrograde (net displacement > 10 µm in the retrograde direction) or bidirectional (net displacement < 10 µm, but total displacement > 10 µm). AVs with net and total displacement < 10 µm were scored as stationary. Non-motile vesicles were binned into bidirectional and stationary vesicles (net and total displacement < 10 mm). A pause was defined as a single or consecutive instantaneous velocity value of < 0.083 µm/sec. Stationary AVs were excluded from the quantification of pause number, pause duration, directional reversals, and Δ run length. Quantification of the fraction of time paused included all (motile and stationary) AVs. For quantification of Δ run length, the net run length of each vesicle was subtracted from its total run length [12]. All analyses were performed by a blinded investigator.

#### Fraction of aSyn-EGFP-positive AVs and number of aSyn-EGFP puncta

Time lapse series were recorded in the proximal axon (< 150 µm from the cell soma) and distal axon (region immediately proximal to the axon tip). Kymographs were generated using the Multiple Kymograph plugin for FIJI as described above. For quantification of the fraction of aSyn-EGFP-positive AVs, traces were counted first in the aSyn-EGFP kymograph, then in the mScarlet-LC3 kymograph by a blinded investigator. For quantification of the number of aSyn-EGFP puncta, the number of aSyn-EGFP-positive puncta present in the first frame of the time lapse acquisition was counted and normalized to 100 µm of axonal length.

#### PFF assay

To quantify the area of pS129 aSyn and MAP2 signal, max projections of each channel were made for each image. Both pS129 aSyn and MAP2 signal were thresholded and binarized in an unbiased manner using ilastik segmentation, a machine-learning based approach to image segmentation [52]. The binarized images were used to quantify pS129-positive area, pS129-positive aggregate number and size, and MAP2 area using the measure function in FIJI.

### Presynapse number

To quantify the number of presynapses (defined as the number of synapsin puncta overlapping with MAP2 signal) in iNeurons and primary mouse cortical neurons, max projections of the synapsin and MAP2 channels were created and thresholded using ilastik segmentation. Using the “AND” function of the image calculator in FIJI, the binarized images were next processed to show only synapsin signal overlapping with MAP2 signal. The analyze particle function (size: 10 pixel^2^ – infinity) and the measure function in FIJI were then used to quantify the number of presynaptic puncta and MAP2 area, respectively.

#### Tagged PFF assay

To quantify intracellular uptake of ATTO 488-labeled PFFs, max projections of each channel were generated. Somata of the imaged cells were outlined manually to create regions of interest by a blinded investigator. ATTO 488 signal was thresholded and binarized using ilastik. Binary images and max projections were overlayed with the created regions of interest and analyzed with the FIJI measure tool to analyze the ATTO 488-positive area and the integrated density, respectively.

#### Statistical analysis

Figure legends contain descriptions of the statistical test(s) used, specific p-values, sample size, and dispersion/precision measurements. For AV transport experiments, statistical analysis of AV directionality was performed by two-way ANOVA with Sidak’s test for multiple comparisons using GraphPad Prism 10. For other transport parameters, RStudio version 2022.07.0 was used to perform a generalized linear mixed model (GLMM; R package ‘‘lme4’’). The genotype (or, in MLi-2 experiments, the treatment condition) was treated as the fixed effect. The independent experiment/culture and the neuron being recorded from were treated as nested random effects, with the neuron nested within the experiment. Specific models were selected as previously described, based on the distribution of each dataset [12]. In detail, the following models were used: for fraction of time paused, GLMM (binomial family, with the ‘‘weights’’ argument for total time); for pause number, GLMM (Poisson family); for pause duration, GLMM (gamma family, log link function); for reversals, GLMM (Poisson family); for Δ run length, GLMM (gamma family, log link function, with transformation to remove zero values with added constants). Statistical analysis of the fraction of AVs positive for aSyn-EGFP, number of aSyn-EGFP positive puncta, pS129 area, MAP2 area, pS129 aggregate size, pS129 aggregate number, as well as area and integrated density of internalized ATTO 488-labeled PFFs was performed by a linear mixed effects model (LME; R package “nlme”) in RStudio. The genotype was treated as the fixed effect. For experiments in primary mouse cortical neurons, the independent experiment/culture and the individual embryo from which neurons were obtained were treated as nested random effects, with the embryo nested within the experiment. For experiments in iPSC-derived iNeurons, the independent experiment/culture was treated as the random effect. For all quantifications, at least three independent experiments were analyzed.

## ACKNOWLEDGEMENTS

We thank the Live-Cell Imaging Facility of the Max Planck Institute for Multidisciplinary Sciences, Goettingen, Germany for providing access to the Visitron CSU-W1 Spinning Disk confocal system and especially Peter Lenart and Antonio Z. Politi for technical support. We thank T. Outeiro, University Medical Center Goettingen, for generously providing ATTO 488-labeled PFFs and Dan Dou, University of Pennsylvania, for helpful insights and discussions. Cartoon schematics were created in part with <u>BioRender.com</u>. The authors gratefully acknowledge support from the Michael J. Fox Foundation (MJFF-021130 to C.A.B. and E.L.F.H.) and the Else Kröner-Fresenius-Stiftung (2023_EKEA.91 to C.A.B.). M.F. and A.M. were supported by intramural fellowships of the University Medical Center Göttingen.

## Abbreviations

aSyn: alpha-synuclein
AV: autophagic vesicle
DIV: day in vitro
iPSC: induced pluripotent stem cell
LRRK2: leucine rich repeat kinase 2
PD: Parkinson disease
PPM1H: protein phosphatase 1H.

