## Supplemental Information for "Knockout of the LRRK2-counteracting RAB phosphatase PPM1H disrupts axonal autophagy and exacerbates alpha-synuclein aggregation"

### Figure S1

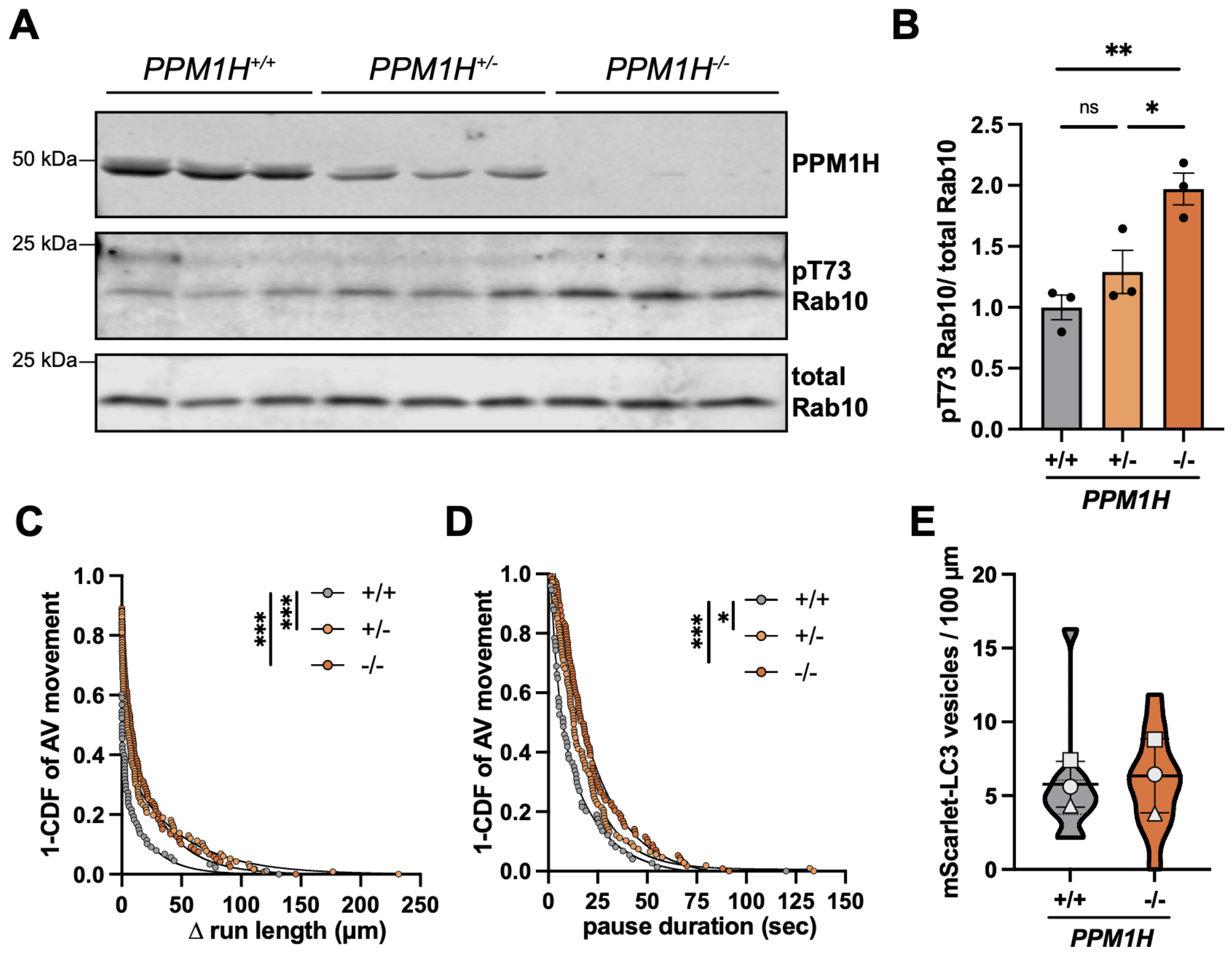


**Figure S1. PPM1H knockout mouse cortical neurons have increased levels of LRRK2-phosphorylated RAB10.** (**A**) Representative Western blot of PPM1H, pT73 Rab10, and total Rab10 in DIV8 *PPM1H*$^{+/+}$, *PPM1H*$^{+/-}$, and *PPM1H*$^{-/-}$ mouse cortical neurons. (**B**) Quantification of pT73 Rab10 normalized to total Rab10 (mean ± SEM; n = 3 biological replicates; ns, not significant, p=0.3669; *p=0.0318; **p=0.0064; One-way ANOVA with Tukey's multiple comparisons test). (**C-D**) Δ run length (C) and pause duration (D) of motile AVs in *PPM1H*$^{+/+}$, *PPM1H*$^{+/-}$, and *PPM1H*$^{-/-}$ mouse cortical neurons (n = 88-119 motile AVs from 27-29 neurons from 3 independent experiments; *p=0.0105; ***p<0.001; mixed effects model analysis, see Methods for specific models used). (**E**) Number of mScarlet-LC3 puncta normalized to 100 µm axonal length in the distal axon of *PPM1H*$^{+/+}$ and *PPM1H*$^{-/-}$ neurons (mean ± SD; n = 11 neurons from 3 independent experiments; ns, not significant, p=0.7897; mixed effects model analysis). For panels C and D, curve fits were generated using nonlinear regression (two phase decay). For panel E, scatter plot points indicate the means of three independent experiments.

### Figure S2

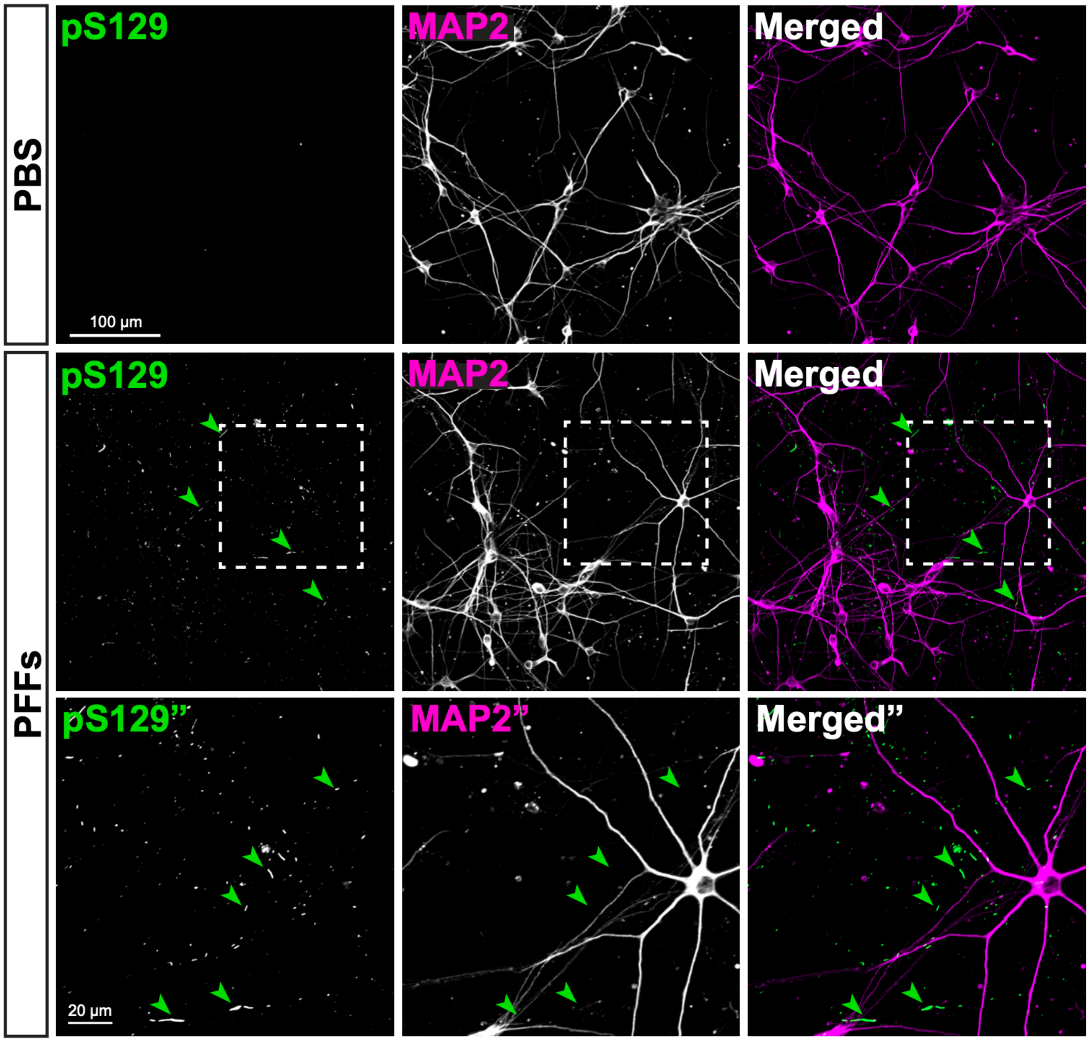


**Figure S2. Exposure to PFFs induces formation of pS129-positive aSyn aggregates in WT iNeurons.** Representative images of DIV35 WT iNeurons stained for pS129-positive aSyn aggregates and MAP2. Cells were exposed to PBS control or 0.25 µM PFFs on DIV14 and fixed on DIV35. Green arrowheads highlight examples of pS129-positive aSyn aggregates. Dashed boxes indicate areas that are shown at higher magnification in the bottom row.

### Figure S3

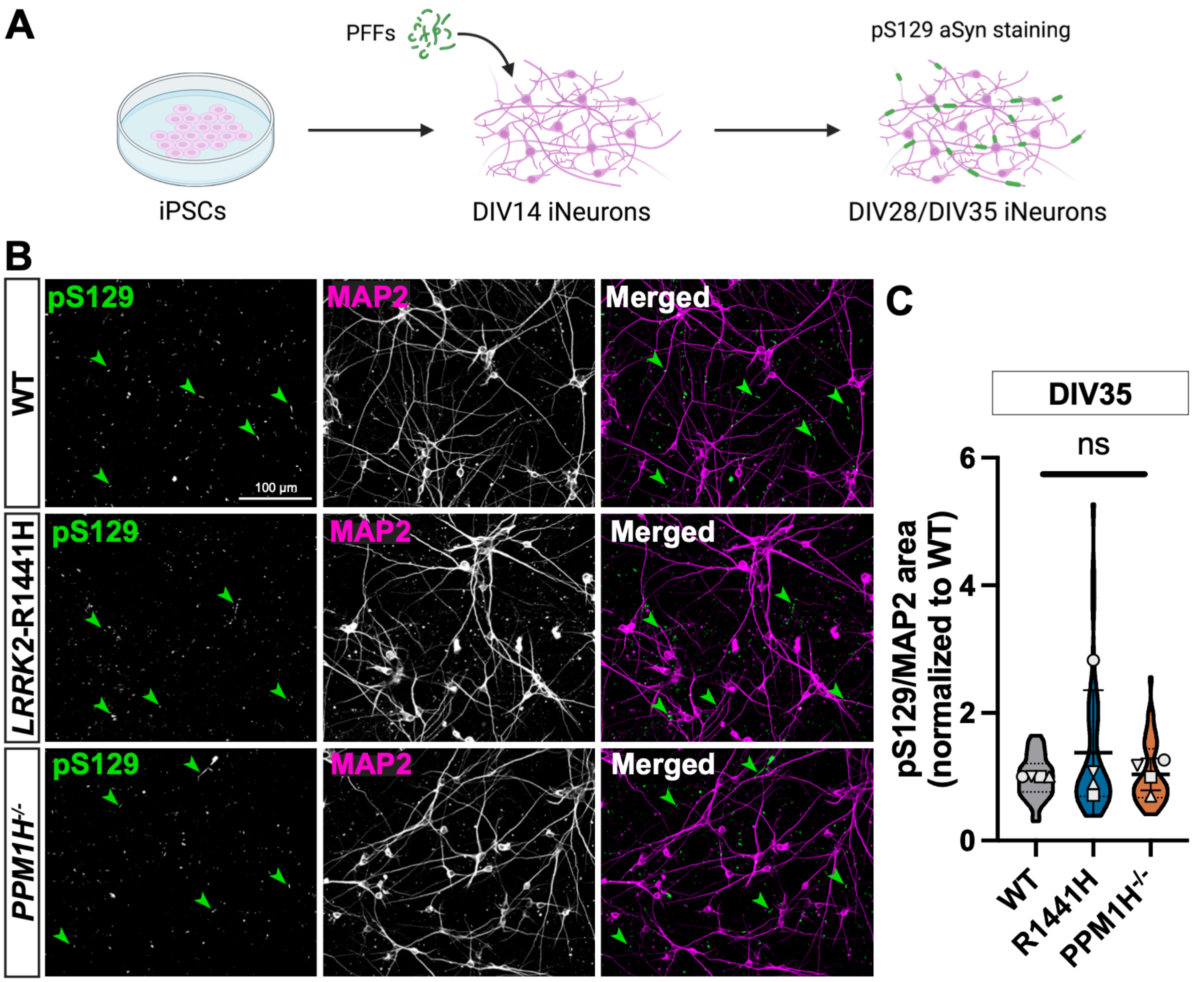


**Figure S3. PFF-induced aSyn aggregation is not altered in PPM1H knockout or *LRRK2*-p.R1441H knockin iNeurons.** (**A**) Schematic depicting the experimental timeline. WT, *LRRK2*-p.R1441H KI, and *PPM1H*$^{-/-}$ iNeurons were exposed to 0.5 µM or 0.25 µM human PFFs on DIV14. iNeurons treated with 0.5 µM PFFs were fixed on DIV28 and iNeurons treated with 0.25 µM PFFs were fixed on DIV35. Fixed neurons were stained for pS129 aSyn and MAP2. (**B**) Representative images of DIV35 WT, *LRRK2*-p.R1441H KI, and *PPM1H*$^{-/-}$ iNeurons stained for pS129-positive aSyn aggregates and MAP2 after 21 days of incubation with PFFs. Green arrowheads highlight examples of pS129-positive aSyn aggregates. (**C**) pS129/MAP2 area normalized to WT control in DIV35 WT, *LRRK2*-p.R1441H KI, and *PPM1H*$^{-/-}$ iNeurons (mean ± SD; n = 40-43 fields of view from 4 independent experiments; ns, not significant, p>0.3551; mixed effects model analysis). For panels C and E, scatter plot points indicate the means of three independent experiments.

### Figure S4

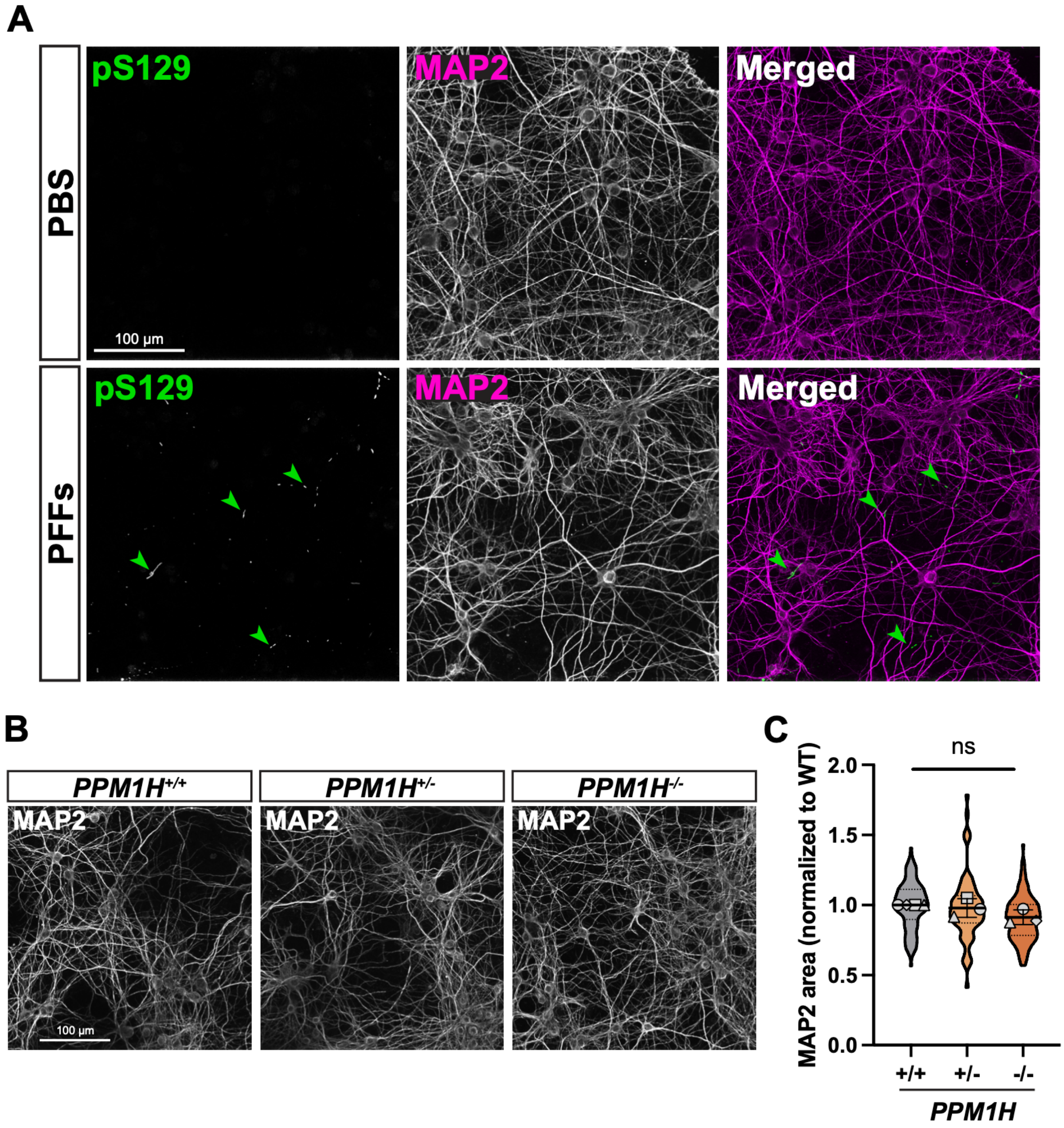


**Figure S4. Exposure to PFFs induces formation of pS129-positive aSyn aggregates and does not result in differences of the MAP2-positive somatodendritic area between *PPM1H*$^{+/+}$, *PPM1H*$^{+/-}$, and *PPM1H*$^{-/-}$ mouse cortical neurons.** (**A**) Representative images of DIV28 WT mouse cortical neurons stained for pS129-positive aSyn aggregates and MAP2. Cells were exposed to PBS control or 0.5 µM PFFs on DIV7 and fixed on DIV28. Green arrowheads highlight examples of pS129-positive aSyn aggregates. (**B**) Representative images of MAP2 staining in DIV28 *PPM1H*$^{+/+}$, *PPM1H*$^{+/-}$, and *PPM1H*$^{-/-}$ mouse cortical neurons treated with 0.5 µM PFFs on DIV7. The same MAP2 images are shown in Figure 3B. (**C**)

Quantification of MAP2 area normalized to WT in DIV28 *PPM1H*$^{+/+}$, *PPM1H*$^{+/-}$, and *PPM1H*$^{-/-}$ mouse cortical neurons treated with 0.5 µM PFFs on DIV7 (mean ± SD; n = 67-120 fields of view from 4 independent experiments; ns, not significant, p>0.0985; mixed effects model analysis). For panel D, scatter plot points indicate the means of each independent experiment.

### Figure S5

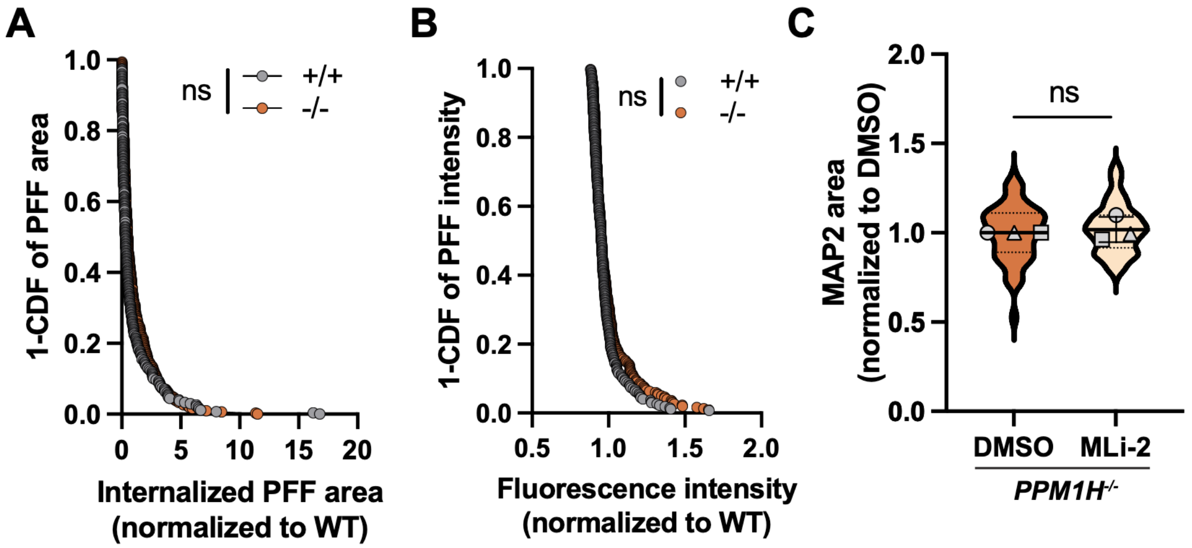


**Figure S5. Knockout of PPM1H does not alter intracellular uptake of fluorescently labeled PFFs.** (**A**) Area of internalized ATTO 488-labeled PFFs normalized to WT control in *PPM1H*$^{+/+}$ and *PPM1H*$^{-/-}$ neurons (n = 270 – 301 neurons from 3 independent experiments; ns, not significant, p=0.3882; mixed effects model analysis). (**B**) Fluorescence intensity (mean gray value) of ATTO 488-labeled PFFs in the soma of DIV8 *PPM1H*$^{+/+}$ and *PPM1H*$^{-/-}$ mouse cortical neurons treated with fluorescently labeled PFFs on DIV7 (n = 270 – 301 neurons from 3 independent experiments; ns, not significant, p=0.6641; mixed effects model analysis). (**C**) Quantification of MAP2 area normalized to DMSO in DIV21 *PPM1H*$^{-/-}$ mouse cortical neurons treated with DMSO or 300 nM MLi-2 from DIV5 until fixation on DIV21 and exposed to PFFs on DIV7 (mean ± SD; n = 98 fields of view from 3 independent experiments; ns, not significant, p=0.5202; mixed effects model analysis). For panel C, scatter plot points indicate the means of three independent experiments.
